# Label-free bone marrow white blood cell classification using refractive index tomograms and deep learning

**DOI:** 10.1101/2020.11.13.381244

**Authors:** DongHun Ryu, Jinho Kim, Daejin Lim, Hyun-Seok Min, Inyoung You, Duck Cho, YongKeun Park

## Abstract

In this study, we report a label-free bone marrow white blood cell classification framework that captures the three-dimensional (3D) refractive index (RI) distributions of individual cells and analyzes with deep learning. Without using labeling or staining processes, 3D RI distributions of individual white blood cells were exploited for accurate profiling of their subtypes. Powered by deep learning, our method used the high-dimensional information of the WBC RI tomogram voxels and achieved high accuracy. The results show >99 % accuracy for the binary classification of myeloids and lymphoids and >96 % accuracy for the four-type classification of B, T lymphocytes, monocytes, and myelocytes. Furthermore, the feature learning of our approach was visualized via an unsupervised dimension reduction technique. We envision that this framework can be integrated into existing workflows for blood cell investigation, thereby providing cost-effective and rapid diagnosis of hematologic malignancy.

## 1. Introduction

Accurate blood cell identification and characterization play an integral role in the screening and diagnosis of various diseases, including sepsis (Baskurt et al. 1998; Bateman et al. 2017; Chandramohanadas et al. 2011; Murphy and Weiner 2012), immune system disorders (Ueda et al. 2003; von Boehmer and Melchers 2010) and blood cancer (Sant et al. 2010). While patients’ blood is examined with regard to the morphological, immunophenotypic, and cytogenetic aspects for diagnosing such diseases (Norris and Stone 2017), the simplest yet the most effective inspection in the early stages of diagnosis is a microscopic examination of stained blood smears obtained from peripheral blood or bone marrow aspirates. In a standard workflow, medical professionals create a blood-smeared slide, fix, and stain the slide with chemical agents su*c*h as Hematoxylin-Eosin and Wright-Giemsa stains, followed by the careful observation of blood cell alternations and cell count as per specific diseases. This not only requires time, labor, and associated costs but also is vulnerable to the variability of staining quality that depends on the staining of trained personnel.

To address this issue, several label-free techniques for identifying blood cells have recently been explored, including multiphoton excitation microscopy (Kim et al. 2018; Li et al. 2010), Raman microscopy (Nitta et al. 2020; Orringer et al. 2017; Ramoji et al. 2012) and hyperspectral imaging (Ojaghi et al. 2020; Verebes et al. 2013). Each method exploits the endogenous contrast (e.g., Tryptophan, Raman spectra, chromophores) of a specimen with the objective of visualizing and characterizing it without using exogenous agents; however, these modalities require rather complex optical instruments with demanding system alignments and long data acquisition time. More recently, quantitative phase imaging (QPI) technologies that enable relatively simple and rapid visualization of biological samples (Bettenworth et al. 2018; Chowdhury et al. 2019; Jung et al. 2018; Kastl et al. 2017; Kim et al. 2014; Kujawińska et al. 2014; Park et al. 2018) have been utilized for various hematologic applications (Chhaniwal et al. 2012; Javidi et al. 2018; Kim et al. 2013; Lee et al. 2017). By measuring the optical path length delay induced by a specimen and by reconstructing a refractive index using the analytic relation between the scattered light and sample, QPI can identify and characterize the morphological and biochemical properties of various blood cells.

Recent advances in artificial intelligence (AI) have suggested unexplored domains of QPI beyond simply characterizing biological samples. As datasets obtained from QPI do not rely on the variability of staining quality, various machine learning and deep learning approaches can exploit uniform-quality and high-dimensional datasets to perform label-free image classification (Chen et al. 2016; Jo et al. 2017; Nissim et al. 2020; Ozaki et al. 2019; Wang et al. 2020; Wu et al. 2020; Yoon et al. 2017; Zhang et al. 2020; Zhou et al. 2020) and inference (Chang et al. 2020; Choi et al. 2019; Dardikman-Yoffe et al. 2020; Kandel et al. 2020; Lee et al. 2019; Nguyen et al. 2018; Nygate et al. 2020; Pitkäaho et al. 2019; Rivenson et al. 2018). Such synergetic approaches for label-free blood cell identification have also been demonstrated, which are of interest to this work (Go et al. 2018; Kim et al. 2019; Nassar et al. 2019; Ozaki et al. 2019; Singh et al. 2020; Yoon et al. 2017). However, these often necessitate manual extraction of features for machine learning or do not fully utilize the high-complexity data of three-dimensional (3D) QPI, possibly improving the performance of deep learning.

In this study, we leverage optical diffraction tomography (ODT), a 3D QPI technique, and a deep neural network to develop a label-free white-blood-cell profiling framework (Fig. 1). We utilized ODT to measure the 3D refractive index of a cell, which is an intrinsic physical property, and extract various morphological/biochemical parameters of the RI tomograms such as cellular volume, dry mass, and protein density. Subsequently, we use the optimized deep learning algorithm for accurately classifying WBCs obtained from bond marrow. To test our method, we performed two classification tasks for a binary differential (myeloid and lymphoid, >99 % accuracy) and a four-group differential (monocyte, myelocyte, B and T lymphocyte, >96% accuracy). We demonstrate the representation learning capability of our algorithm using unsupervised dimension reduction visualization. We also compare conventional machine learning approaches and two-dimensional (2D) deep learning with this work, verifying the superior performance of our 3D deep learning approach. We envision that this label-free framework can be extended to a variety of subtype classifications and has the potential to advance hematologic malignancy diagnosis.

**Fig. 1:**
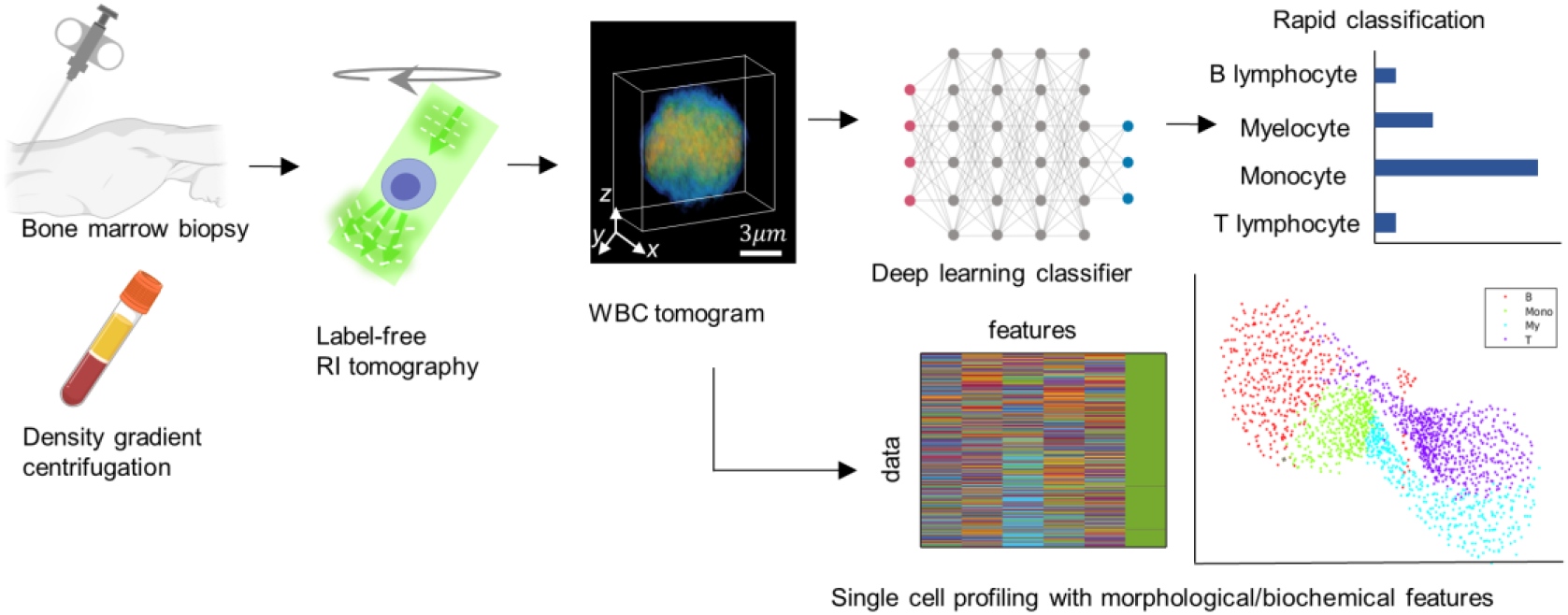
Overview of label-free bone marrow white blood cell (WBC) assessment using diffraction tomography and artificial intelligence. Bone marrow WBCs obtained from a minimal process of density gradient centrifugation are tomographically imaged without any labeling agents. Subsequently, individual refractive index (RI) tomograms can be accurately (>95%) and rapidly (<1 s) classified via an optimized deep-learning classifier along with the analysis of morphological/biochemical features such as cellular volume, dry mass, and protein density.

## 2. Material and methods

### 2.1. Optical diffraction tomography

Our 3D imaging system is a commercialized optical diffraction tomography microscope (also known as holotomography) using Mach–Zehnder interferometry and a digital micromirror device (DMD) for high-speed angle-scanned illumination (HT-2H, Tomocube, Inc., South Korea) (Shin et al. 2016). The schematics of its optical setup are depicted in Fig. S1 (a) (Refer to Supplementary file). A diode-pumped solid-state laser beam (532-nm wavelength, MSL-S-532-10mW, CNI laser, China) was used for the sample beam and reference beam, split by a fiber coupler (OZ optics, Canada). The sample beam, angle-scanned by a DMD (DLP65300FYE, Texas Instruments, USA), impinges on a sample after passing through a condenser objective lens (UPLASAPO 60XW, Olympus Inc., Japan). The scattered signal, 4-*f*-relayed by an objective lens (UPLASAPO 60XW, Olympus Inc., Japan) and a tube lens, interferes with the reference beam transmitted via a beam splitter. After being filtered by a linear polarizer, 49 interferograms were captured by a complementary metal-oxide-semiconductor camera (FL3-U3-13Y3M-C, FLIR Systems, Inc., USA), as shown in Fig. S1 (b).

Next, the amplitude and phase images are retrieved from the measured interferograms using a phase-retrieval algorithm that utilizes spatial filtering (Cuche et al. 2000) and the Goldstein phase unwrapping method (Goldstein et al. 1988). Based on the Fourier diffraction theorem with Rytov approximation (Wolf 1969), a sample’s 3D RI distribution is reconstructed from the retrieved amplitude and phase images. To fill in the side-scattering signal not collected owing to the limited numerical apertures of objective lenses, a regularization algorithm utilizing a non-negative constraint is employed (Sung and Dasari 2011). The theoretical resolutions of the ODT system are 110 nm (lateral) and 330 nm (axial), according to the Lauer criterion (Lauer 2002). The data acquisition time for the 49 interferograms is approximately 500 ms, and it requires typically several seconds to reconstruct a regularized tomogram using a standard personal computer. The reconstruction time for the tomogram can be further reduced by a high-end graphics processing unit. All these customized image reconstruction codes were implemented in MATLAB (MathWorks, USA).

### 2.2. Sample preparation and data acquisition

Four types of white blood cells were obtained from the bone marrow of eleven healthy donors investigated: myelocyte, monocyte, B lymphocyte, and T lymphocyte (Fig. 2(a)). In WBC lineage, myelocytes and monocytes stem from myeloids; B and T lymphocytes stem from lymphoids. The study was approved by the Institutional Review Board (No. SMC 2018-12-101-002) at the Samsung Medical Center (SMC). All blood samples were acquired at the SMC between November 2019 and June 2020. We selected patients suspected of having lymphoma who underwent bone marrow biopsy and were all diagnosed with a normal bone marrow at the end.

**Fig. 2:**
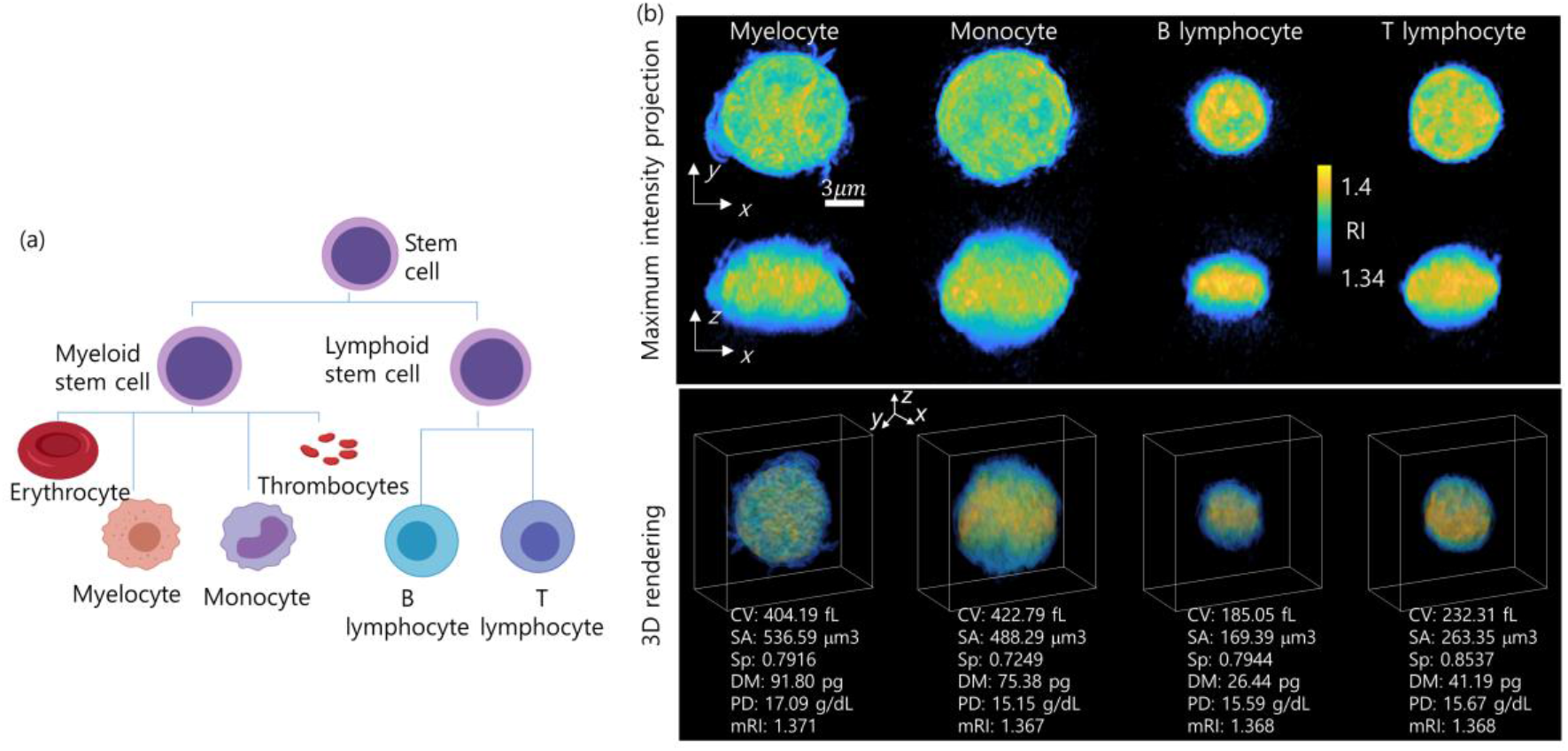
Investigated WBC lineage and representative RI tomograms. (a) Two myeloid cells and two lymphoid cells are used to demonstrate our framework. (b) Representative RI tomograms are visualized through two different perspectives of maximum intensity projection and 3D isosurface rendering. CV: Cellular Volume. SA: Surface Area. Sp: Sphericity. DM: Dry Mass. PD: Protein Density. mRI: mean RI.

First, the bone marrow was extracted via a needle biopsy. To isolate mononuclear cells (MC), the bone marrow was diluted in phosphate-buffered saline (PBS; Welgene, Gyeongsan-si, Gyeongsangbuk-do, Korea) in a 1:1 ratio, centrifuged using Ficoll-Hypaque (d = 1.077, LymphoprepTM; Axis-Shield, Oslo, Norway), and washed twice with PBS. Next, magnetic-activated cell sorting was performed to obtain the four types of WBCs from the isolated MC. CD3+, CD19+, and CD14+ MACS microbeads (Miltenyi Biotec, Germany) were used to positively select T, B, and monocytes. Myelocyte cells were isolated through the negative selection of CD14 and positive selection of CD33. For optimal sample density and viability, we prepared each isolated sample in a mixed solution of 80% Roswell Park Memorial Institute (RPMI) 1640 medium, 10% heat-inactivated fetal bovine serum (FBS), and 10% 100 U/mL penicillin and 100-µg/mL streptomycin (Lonza, Walkersville, MD, USA). While imaging, we kept the samples in an enclosed iced box.

A total of 2547 WBC tomograms were obtained using our imaging system. The number of datasets for myelocytes, monocytes, B lymphocytes, and T lymphocytes was 610, 516, 508, and 913, respectively. For deep learning, we randomly shuffled the entire dataset and split the training, validation, and test set by a 7:1:2 ratio.

The representative 3D RI tomograms for each subtype are visualized as a maximum intensity projection image and a 3D rendered image in Fig. 2(b). The morphological and biochemical parameters can be directly computed from the measured RI tomograms, which is further explained in the next section.

### 2.3. Quantitative analysis of morphological/biochemical properties

Six parameters calculated from a reconstructed tomogram were obtained: cellular volume, surface area, sphericity, dry mass, protein density, and mean refractive index. First, the cellular volume and surface area were directly acquired by thresholding the tomogram. The voxels with RI values higher than the thresholding RI value were segmented; the thresholding value was 1.35, considering a known medium RI of approximately 1.33, and experimental noises. Sphericity was calculated by relating the obtained surface area S and volume V as follows: Sphericity = π^1/3^·(6 V) ^2/3^/S.

Next, the biochemical properties such as protein density and dry mass were obtained from RI values via a linear relation between the RI of a biological sample and its local concentration of non-aqueous molecules (i.e., proteins, lipids, and nucleic acids inside cells). Considering that the proteins are major components and have mostly uniform values, protein density can be directly converted from RI values as follows: *n* = *n*_0_+*αC*, where *n* and *n*_0_ are the RI values of a voxel and the medium, respectively, and α is the refractive index increment (RII). In this study, we used an RII value of 0.2 mL/g. The total dry mass can be calculated by integrating the protein density over the cellular volume.

### 2.4. Conventional machine learning approaches

To compare our deep learning approach with existing machine learning approaches, we implemented support vector machine (SVM), k-nearest neighbors (KNN), linear discriminant classifier (LDC), naïve Bayes (NB), and decision tree (DT) algorithms to classify the four types of WBCs using six extracted features (cellular volume, surface area, sphericity, dry mass, protein density, and mean RI). Six binary-SVM models with error-correcting output codes were trained to make a decision boundary for the four classes. For the KNN classifier, k = 5 was chosen. All machine learning algorithms were implemented using MATLAB.

### 2.5. Deep neural network for the classification of BM WBCs

We implemented a deep neural network to identify the 3D RI tomogram of myelocytes, monocytes, T lymphocytes, and B lymphocytes. Our convolutional neural network, inspired by FISH-Net(Sun et al. 2018), comprises two downsampling (DS) blocks, an upsampling (US) block, and a classifier at the end (Fig. 3). The first DS block, comprising batch norm, leaky ReLu, 3D convolution, and 3D max pooling, extracts various features at low resolution. Next, the US block upsamples using nearest-neighbor interpolation and refines the features obtained from the previous block via residual blocks that connect all three DS and US blocks together. These residual blocks help the improved flow of information across different layers and relax the well-known vanishing gradient. The second DS block processes not only the previous features from the US block but also the features transmitted via the residual blocks with batch norm, leaky ReLu, 3D convolution, and 3D max pooling. Ultimately, the extracted features, after being processed by the classifier comprising batch norm, leaky Relu, 3D convolution, and adaptive 3D pooling, are set to the most probable subtype for the classification task.

**Fig. 3:**
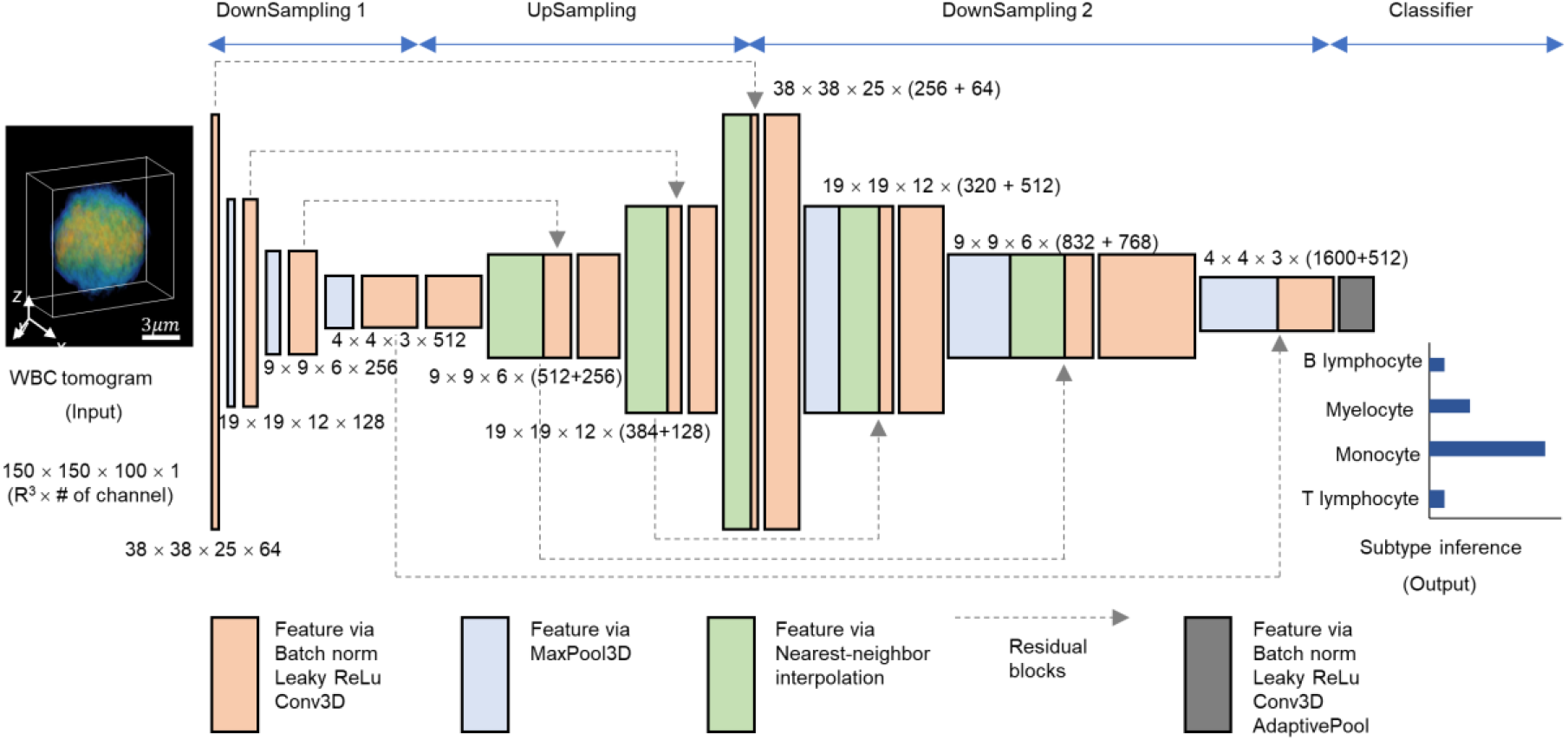
Three-dimensional deep neural network for WBC classification of BM WBCs. RI tomograms are processed to extract distinguishing features of each WBC through a convolutional neural network comprising downsampling and upsampling operations, leading to the subtype inference at the end.

Our network was implemented in PyTorch 1.0 using a GPU server computer (Intel® Xeon® silver 4114 CPU and 8 NVIDIA Tesla P40). We trained our network using an ADAM optimizer (Kingma and Ba 2014) (learning rate = 0.004, momentum = 0.9, and learning rate decay = 0.9999) with a cross-entropy loss. The learnable parameters were initialized by He initialization (He et al. 2015). We augmented the data using random translating, cropping, elastic transformation, and adding Gaussian noise. We trained our algorithm with a batch of 8 for approximately 18 h. We selected our best model at 159 epochs and monitored the validation accuracy based on MSE.

### 2.6. Uniform manifold approximation and projection (UMAP) for dimension reduction

To effectively visualize the learning capability of our deep neural network, we employed a cutting-edge unsupervised dimension reduction technique known as uniform manifold approximation and projection (UMAP) (McInnes et al. 2018). UMAP constructs the high-dimensional topological representations of the entire dataset and optimizes them into low-dimensional space (e.g., two dimensions) to be as topologically similar as possible. We extracted the learned features from the last layer of the second downsampling block immediately before entering the classifier block and applied UMAP to demonstrate the feature learning capability of the trained classifier in two dimensions. We used open-sourced UMAP implemented in MATLAB (Meehan et al. 2020).

## 3. Results and discussion

### 3.1. Morphological and biochemical properties of WBCs

We first quantified morphological (cell volume, surface area, and sphericity), biochemical (dry mass and protein density), and physical (mean RI) parameters of bone marrow WBCs and statistically compared them in Fig. 4. These parameters can be directly obtained from the 3D RI tomograms (refer to section 2.3). The mean and standard deviation of cell volumes for B lymphocytes, monocytes, myelocytes, and T lymphocytes are 185.05 ± 34.86 fL, 422.79 ± 61.31 fL, 404.19 ± 53.10 fL, and 232.31 ± 30.35 fL, respectively; in the same order, the surface areas are 169.39 ± 49.21 μm^3^, 498.29 ± 80.60 μm^3^, 536.59 ± 91.01 μm^3^, and 263.35 ± 48.84 μm^3^; the sphericities are 0.794 ± 0.026, 0.725 ± 0.076, 0.792 ± 0.057, and 0.854 ± 0.022; the dry masses are 26.44 ± 8.13 pg, 75.38 ± 14.01 pg, 91.80 ± 17.51 pg, and 41.19 ± 7.88; the protein densities are 15.58 ± 1.05 g/dL, 15.15 ± 1.55 g/dL, 17.09 ± 1.26 g/dL, and 15.67 ± 0.83 g/dL; the mean RIs are 1.368 ± 0.002, 1.367 ± 0.003, 1.371 ± 0.002, and 1.368 ± 0.001.

**Fig. 4:**
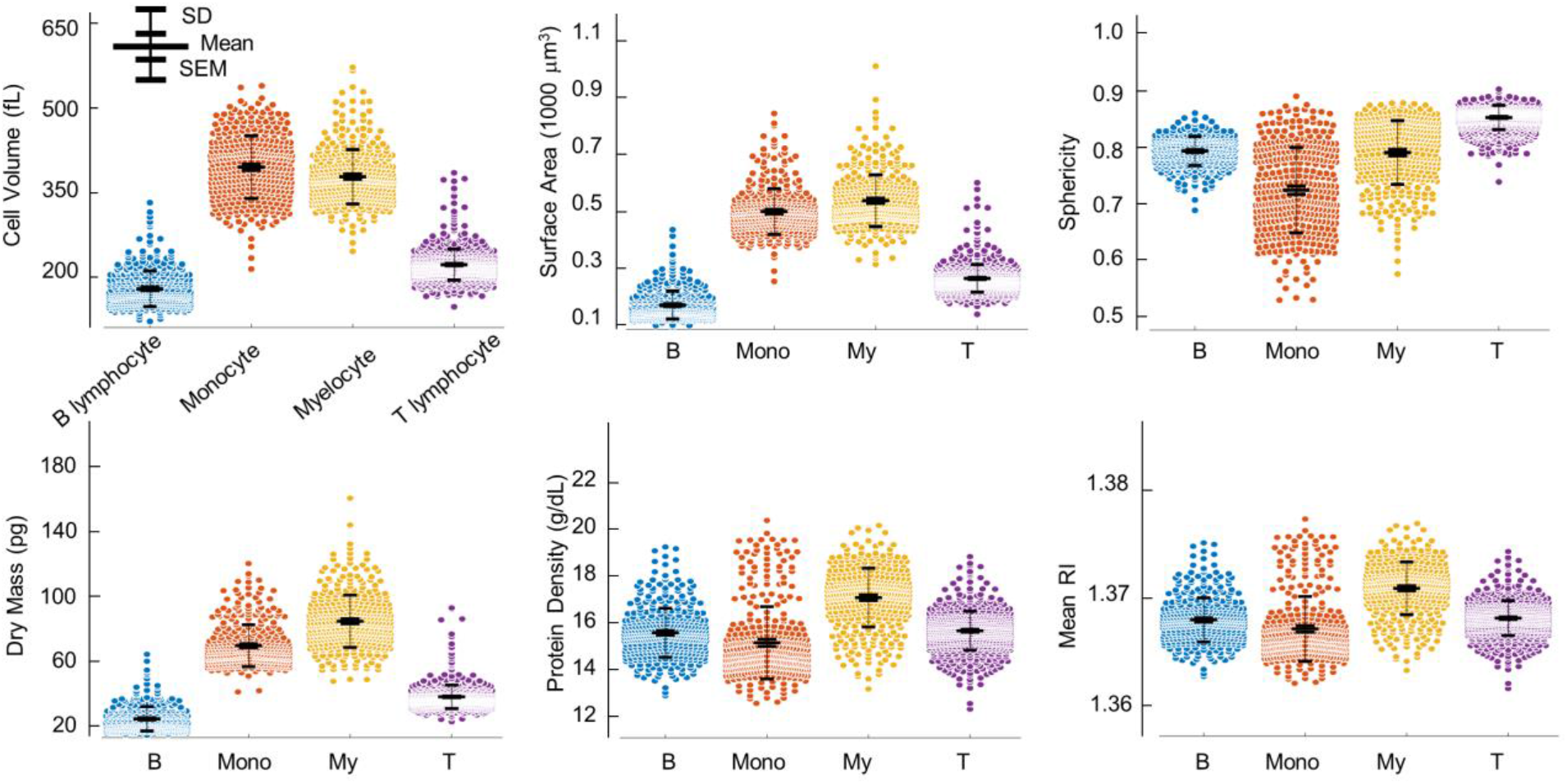
Quantitative analysis of morphological (cell volume, surface area, and sphericity), biochemical (dry mass and protein density), and physical (mean RI) parameters. The protein density is directly related to the mean RI. The scatter plots represent the entire population of the measured tomograms. SD: standard deviation. SEM: standard error of the mean. B: B lymphocyte. Mono: monocyte. My: myelocyte. T: T lymphocyte.

Several observations are noteworthy. First, the mean cellular volume, surface, and dry mass of lymphoid cells (B and T lymphocytes) are smaller than those of myeloid cells (monocytes and myelocytes). The morphological properties of the B and T cells are directly related to the dry mass because we assumed that the cells mainly comprised proteins (0.2 mL/g). Furthermore, we observed that the sphericity of the lymphoid group was larger than that of the myeloid group. The B and T lymphocytes, which commonly originate from small lymphocytes, have one nucleus and spherical shapes; the monocyte and myelocyte cells have a more irregular morphological distribution. Overall, the standard deviations of all parameters except the mean RI for the myeloid group were larger than those of the lymphoid group, indicating the larger morphological and biochemical variability in the myeloid group. Finally, it is challenging to accurately classify the four types of WBCs by simply comparing these parameters (e.g., thresholding). Although lymphoid cells could be rather differentiated from myeloid cells based on the cellular volume, surface area, or dry mass at the “population” level, the overlapped population on all parameters across the two groups still impedes the accurate classification at the “single-cell” level. More importantly, classifying within lymphoid or myeloid (e.g., classification of B and T) would be even more difficult as their statistical parameters are very similar.

### 3.2. Three-dimensional deep learning approach for accurate classification

To achieve an accurate classification of the WBCs at the single-cell level, we designed and optimized a deep-learning-based classifier by fully exploiting the ultra-high dimension of WBC RI voxels, presenting two independent classification results in Fig. 5. We first tested the 3D deep-learning classifier for the binary classification of lymphoid and myeloid cell groups. For the unseen test dataset, the binary classification accuracy of the trained algorithm was 99.7 %, as depicted in Fig. 5 (a). Remarkably, only one lymphoid-group cell was misclassified as a myeloid cell. The powerful learning capability of our network is visualized by the unsupervised dimension reduction technique, UMAP (see Section 2.6) in Fig. 5(b). High-dimensional features were extracted from the last layer of the second downsampling block of the trained network. A majority of test data points are clearly clustered, while few data points of myeloids and lymphoids are closely located. This indicates that our well-trained algorithm not only extracts various features that differentiate the two groups but also finely generates a complex decision boundary for such unclustered data points for accurate classification. It is also interesting that roughly four clusters were generated through deep learning, although we trained the algorithm to classify the two groups, implying that the learning capacity of our algorithm would be sufficient for the classification of more diverse subtypes.

**Fig. 5:**
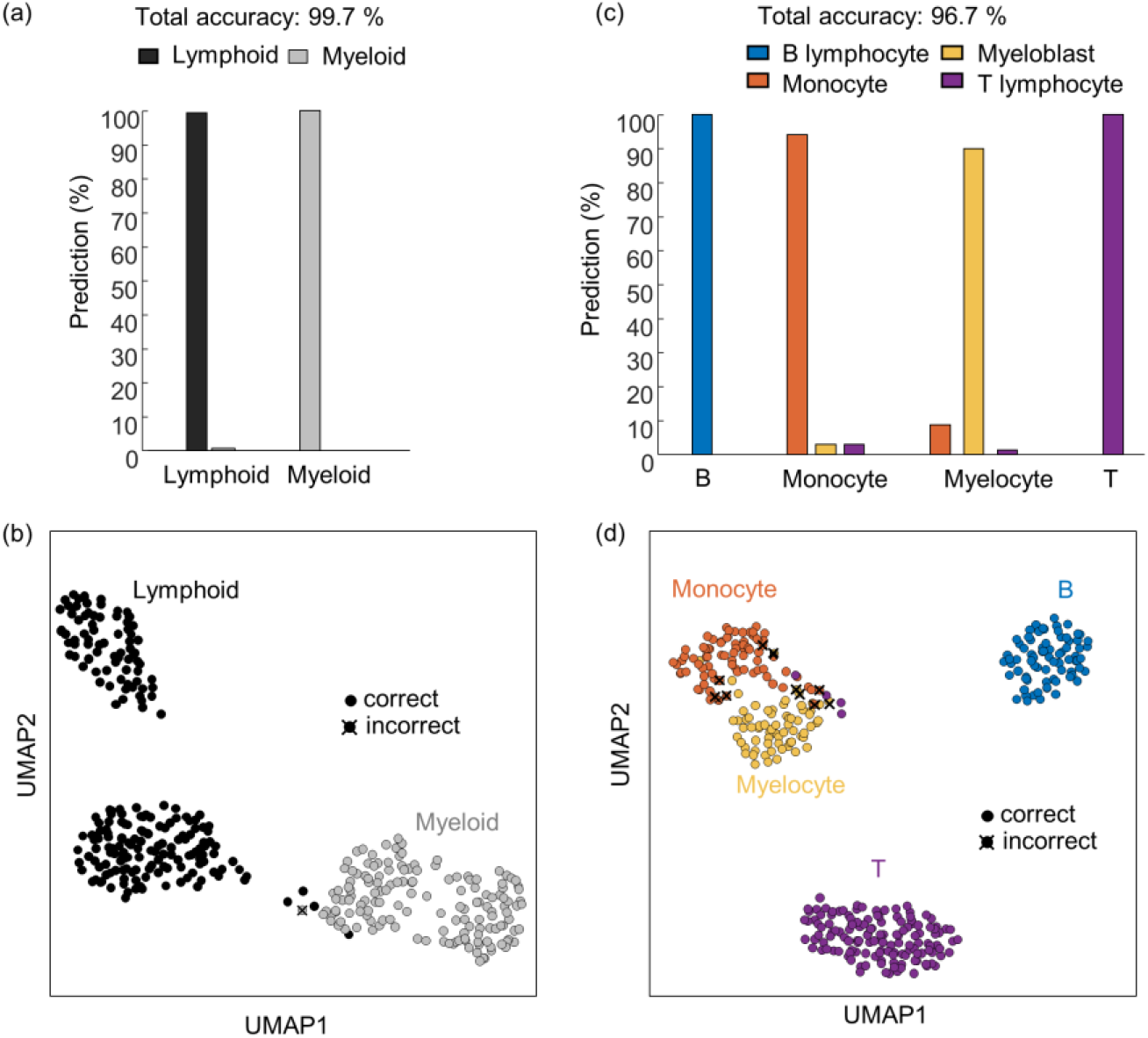
Performance of our optimized deep-learning classifiers. Two classifiers are independently designed for (a, b) the binary classification of lymphoid and myeloid and (c, d) the classification of B/T lymphocytes, monocyte, and myelocyte, respectively. The powerful learning capability of the optimized classifiers is illustrated via uniform manifold approximation and projection (UMAP).

We also tested another deep neural network that classifies the four types of WBCs with a test accuracy of 96.7%. The predictions of the trained algorithm for the four different subtypes are shown in Fig. 5 (c). Our algorithm correctly classified all B lymphocytes, while monocyte and myelocyte groups were misclassified, and a few T lymphocytes were misclassified into myeloid group cells. UMAP visualization was also performed for this result (Fig. 5(d)). The B cell cluster is clearly distant from the remaining clusters. Meanwhile, the monocyte and myelocyte clusters are closely located, thereby providing the most cases of misclassification. A few data points of T cells were also found near these clusters. Despite the significantly similar statistics across the four subtypes as confirmed in the previous section, our algorithm is capable of extracting useful visual features from millions of voxels and generating a sophisticated decision boundary, achieving high test accuracy for the unseen data.

### 3.3. Conventional machine learning and 2D deep learning

For further validation of our method, we benchmarked our 3D deep learning approach against conventional machine learning (ML) approaches that require handcrafted features. First, widely used ML algorithms, such as support vector machines, k-nearest neighbors, linear discriminant classifiers, naïve trees, and decision trees, were performed and compared with our method for the four-type classification (refer to Section 2.4). The test accuracies for the five algorithms along with our method are shown in Fig. 6 (a). While the machine learning algorithms do not exceed 90% accuracy, our method achieved more than 96% test accuracy, as confirmed in the previous section. We reasoned that the six parameters obtained from 3D RI tomograms would not be sufficient for the conventional algorithms to generate an accurate classification boundary of the 3D deep network, although additional feature engineering or extractions may help improve the performance of the machine learning algorithms.

**Fig. 6:**
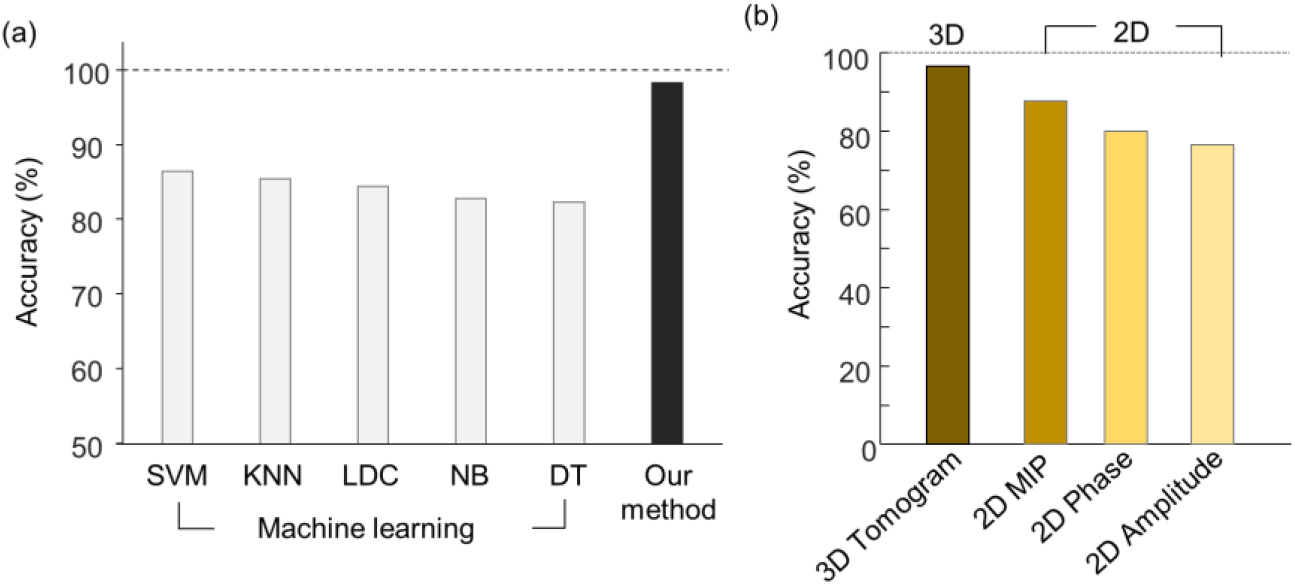
(a) Comparison of classification accuracy between conventional machine learning algorithms that require hand-crafted features and our 3D deep learning. (b) performance of deep learning with various input data. SVM: Support Vector Machine. KNN: K-Nearest Neighbors algorithm. LDC: Linear Discriminant Classifier. NB: Naïve Bayes. DT: Decision Tree. MIP: Maximum Intensity Projection.

A 2D deep neural network that processes various 2D input data was also explored. We used a 2D maximum intensity projection (MIP) image that can be directly obtained from a 3D RI tomogram, 2D phase, and amplitude to train the deep network. The classification accuracies for the four different inputs are shown in Fig. 6 (b). While the 2D MIP obtained directly from the 3D tomogram achieved an accuracy of 87.9%, which is the closest to our approach, the network trained with 2D phase and amplitude presented accuracies of 80.1% and 76.6%. These results suggest that it is important to fully utilize 3D cellular information with an optimal 3D deep-learning classifier for accurate classification.

## Conclusions

In this study, we have demonstrated how synergistically utilizing optical diffraction tomography and deep learning can be employed to profile bone marrow white blood cells. With minimal sample processing (e.g., centrifugation), our computational framework can accurately classify white blood cells (lymphoids and myeloids with >99% accuracy; monocytes, myelocytes, B, and T lymphocytes with >96% accuracy) and their useful properties such as cellular volume, dry mass, and protein density without using any labeling agents. Moreover, the presented approach, capable of extracting optimal features from captured RI voxels, outperformed the well-known machine learning algorithms requiring handcrafted features and deep learning with 2D images. The powerful feature learning capability of our deep neural networks was demonstrated via UMAP, an unsupervised dimension reduction technique. We anticipate that this label-free workflow that does not require laborious sample handlings, and domain knowledge can be integrated into existing blood tests, enabling the cost-effective and faster diagnosis of related hematologic disorders.

Despite successful demonstrations, several future studies need to be conducted for our approach to be applicable in clinical settings. First, diverse generalization tests across imaging devices and medical institutions should be performed. In this study, we acquired the training and test datasets from only one imaging system installed at a single site. Exploring diverse samples and cohort studies at multiple sites using several imagers would strengthen the reliability of our approach. Second, the current data acquisition rate needs to be improved. While we manually obtained and captured the tomograms of WBCs within the limited field-of-view of high numerical aperture (30 × 30 um^2^, 1.2 NA), the use of a motorized sample stage well synchronized with a high-speed camera can significantly boost the data acquisition rate. The microfluidic system might be integrated into the current system to improve the imaging speed; however, tomographically imaging rapidly moving samples as accurately as static ones would be challenging. Ultimately, we need to extend the classification to other types of bone marrow blood cells for diagnosing various hematologic diseases such as leukemia, which may require the additional tuning of our deep network.

## Acknowledgements

We thank Young Seo Kim (KAIST) and Anna Yoon (MIT) for assisting in sample preparation protocol.

